# MEM-GAN: A Pseudo Membrane Generator for Single-cell Imaging in Fluorescent Microscopy

**DOI:** 10.1101/2023.11.08.566343

**Authors:** Yixin Wang, Jiayuan Ding, Lidan Wu, Aster Wardhani, Patrick Danaher, Wenzhuo Tang, Hongzhi Wen, Qiaolin Lu, Yi Chang, Yu Leo Lei, Jiliang Tang, Yuying Xie

## Abstract

Fluorescent microscopy imaging is vital to capturing single-cell spatial data, characterizing tissue organization and facilitating comprehensive analysis of cellular state. Advancements in fluorescent microscopy imaging technologies have enabled precise downstream cellular analysis, particularly in cell segmentation. Accurate segmentation of individual cells allows better profiling and understanding of cell properties and behaviors. The majority of existing segmentation methods predominantly concentrate on enhancing segmentation algorithms, and their effectiveness strongly relies on the input stained image quality. Factors such as high cellular density, indistinct cell boundaries, and staining artifacts can result in uneven and low-quality staining, particularly causing missing or unclear membrane staining. These artifacts adversely impact the efficacy of the subsequent cell segmentation methods. To tackle this insufficient membrane staining, we propose a novel approach, Mem-GAN, to generate high-quality membranes for cells with missing or weak membranes. Inspired by advanced style transfer techniques in computer vision, Mem-GAN styles the content of the cells with missing or weak membranes into cells with integrated membrane staining. Considering the differences in membrane morphology between epithelial/tumor cells and immune cells, Mem-GAN deals with tumor and immune cells separately, not only enhancing membrane staining for cells with partially weak membrane signals but also generating membranes for cells with only nuclear channels. The proposed Mem-GAN is evaluated using the publicly available CosMx dataset. Experimental results demonstrate significant improvements in image staining quality, more accurate representation of membrane morphology characteristics, and better performance in downstream segmentation tasks. Mem-GAN is flexibly adapted and applied to other spatially resolved transcriptomics datasets, such as MERFISH and FISHseq. Our work provides a new perspective on tackling the challenges in cell segmentation from fluorescent microscopy image restoration. The implementation of Mem-GAN is open-source and available at the github repository https://github.com/OmicsML/Mem-GAN. The interactive webserver-based demo of Mem-GAN can be accessed at http://omicsml.ai/memgan.

## 1 INTRODUCTION

Fluorescent microscopy imaging is essential technique to capturing single-cell spatial information, assisting in further analyses and understanding of cellular states and tissue structures. Immunohistochemistry (IHC) [3] has been a widely used multiplexed imaging method since its debut in 1942. IHC uses appropriately labeled antibodies to specifically bind to their target antigens in situ, which can be captured more effectively by current light or fluorescence microscopy. With advances in multiplexed imaging technologies, various techniques [4, 6, 7, 16, 21, 22] have been developed to enhance individual cell resolution. These images are usually acquired using sequential antibody staining and dye as protein markers for cellular identification. These markers effectively label specific cell structures and shapes within a sample and have been continuously improved to enhance sensitivity and specificity for single-cell analysis.

The advances in fluorescent microscopy imaging technologies allow for precise downstream analysis on the cellular level to understand underlying biological processes, such as cell segmentation, cell type recognition, and cell tracking. Single-cell segmentation identifies and separates individual cells within an image by creating pixel-level labels or annotations for each cell. This segmentation enables the study of single cells within a population and allows the profiling and understanding of the properties and behaviors of individual cells in an image-based manner. Single-cell segmentation from fluorescent microscopy imaging, often using ideas from traditional computer vision techniques, has received increased attention. Early works, such as Yan et al.[36] and MxIF [5], utilized pre-selected markers for segmentation, while Schüffier et al. [31] developed an automatic marker selection method to capture multiplexed imaging information. Methods, like SVSS [29] and Matisse [18], incorporated prior knowledge of cell shape and multiple marker information to enhance segmentation performance. These methods leveraged various tools, such as the watershed algorithm, level set methods, and open-source technologies like Ilastik [32] and CellProfiler [15]. Recently, deep learning (DL) has greatly advanced computer vision and image segmentation, where numerous robust algorithms have been proposed to greatly improve cell segmentation performance[9, 17, 23, 26, 33, 34]. The U-Net architecture [27] is a popular Convolutional Neural Network (CNN) used for medical image segmentation tasks, but it is prone to fail in overlapping cells. One solution involves using a regression model that predicts continuous variables for each pixel instead of categorizing them as foreground or background. This approach has been successfully implemented in frameworks such as Cellpose [33, 34] and DeepDistance [17]. In addition, TissueNet [9] is a large dataset that constructs a more specified and robust structure for cell segmentation, and MIRIAM [23] is a pipeline that handles highly multiplexed imaging platforms.

Single-cell segmentation highly depends on the simultaneous staining of cell nuclei and cell membranes. The high staining quality benefit cell segmentation performed by software tools [13]. Typically, cell nuclei are stained using DAPI, a fluorescent dye that binds to DNA, while various membrane protein antibodies like CD45, CD3, and PanCK are employed to label the cell membranes. Despite the advancements in imaging techniques, which involve carefully selecting antibodies and incorporating of additional antibodies for enhanced multiplexing capabilities, introducing noises into multiplexed immunofluorescence images is inevitable. These noises can manifest as cells without membrane staining or cells with weak membrane staining, leading to low-quality and uneven staining, as depicted in Figure. 1 (Original images). These issues can arise from both optical artifacts, such as out-of-focus or blurry images, and chemical artifacts, including variations in antibody concentration and epitope availability in the samples. Furthermore, when employing multi-modality assays that measure both RNA and protein in the same section rather than assays specifically targeting proteins, compromised staining quality with weaker membrane signals can occur.

**Figure 1:**
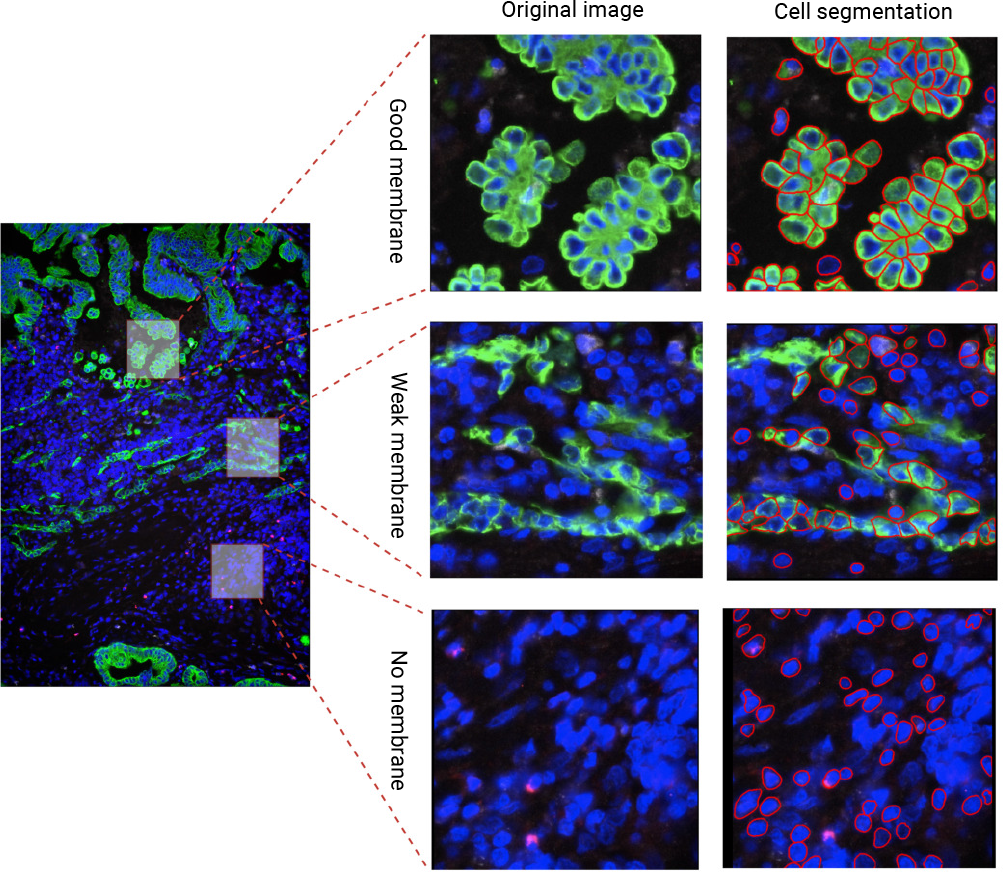
Cells with distinct membrane staining qualities and corresponding cell segmentation results from Cellpose [34] on CosMx dataset (“Lung5 Rep1” sample). Cells with good membrane staining yield highly accurate cell segmentation outcomes, while cells without membrane staining or with weak membrane staining result in comparatively poorer results.

Moreover, during the pre-processing process of the raw images stained with different membrane biomarkers, carefully choosing different antibodies and well-designed aggregation strategies for these images ensures the maximum detection of the target and the appropriate definition of the cell boundaries [1]. Inappropriate choices exacerbate the difficulty of obtaining high-quality images with high cell border contrast. Therefore, even the most sophisticated segmentation methods heavily rely on the quality of the provided stained images and are prone to failure when faced with instinct cell boundaries resulting from weak or missing membrane staining. As depicted in Figure. 1, the widely used segmentation tool Cellpose [34] accurately delineates the boundaries of individual cells. However, it struggles when cells exhibit weak or no membrane staining, which highlights the challenge of incomplete or faint membrane signals.

Most existing works on single-cell segmentation focus on improving the algorithms, but overlook the vital role of the fluorescence quality plays. To handle cells with weak cell membrane staining, most methods directly segment the cell nuclei, while others enlarge the region from the nuclei. Schmitt et al. [30] performs a morphological reconstruction through dilation and erosion operations and Lin et al. [22] utilizes Voronoi tessellation. To handle more irregular and elongated cell shapes, McKinley et al. [24] finds the endpoints of internal membranes and extends them until they intersect with other cells. However, these methods are to refine the segmentation results, which could fail to characterize the real morphologies of different cell membranes.

Rather than merely extending segmentation boundaries, we take a novel perspective that generates superior cell membranes for cells with missing or weak membranes. Drawing inspiration from the advanced style transfer technique in the computer vision domain, Cycle-consistent Generative Adversarial Network (CycleGAN) [39], we develop a membrane generator called Mem-GAN. This generator styles the content of cells with missing or weak membranes into cells with well-defined membrane staining. Mem-GAN effectively generates improved cell membranes, enhancing image staining quality and capturing more accurate membrane morphology characteristics while preserving the cellular structure.

Membrane generation models should also consider the exhibited membrane differences between different cell types. As shown in Figure. 2, tumor cells often display irregular and heterogeneous shapes, resulting in a more jagged or uneven membrane appearance. In contrast, immune cells, such as lymphocytes, typically have a more rounded and uniform shape, characterized by a smooth and continuous membrane. Therefore, Mem-GAN is also trained to deal with tumor and immune cells separately. Since, in the most cases, cells are often losing part of the staining signals, Mem-GAN either generates membranes for cells with only nuclei channels or enhances membranes for cells with partially weak membrane signals. The results can be qualitatively and quantitatively analyzed in the context of generated membrane accuracy and downstream segmentation evaluation. The experiments are conducted on the publicly released dataset, CosMx [10], and Mem-GAN can be easily adapted to other spatially resolved transcriptomics datasets, such as MERFISH and FISHseq.

**Figure 2:**
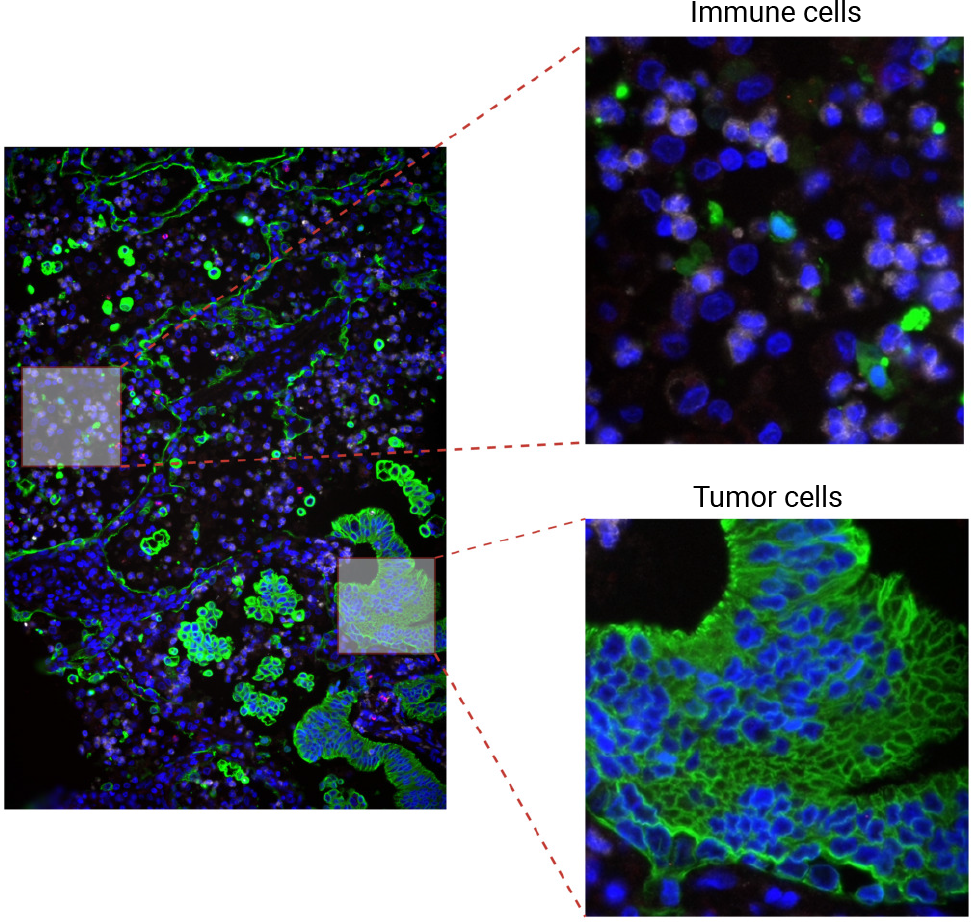
A comparison between immune cells and tumor cells highlights distinctions in their membrane shapes and fluorescent staining signals.

## 2 MATERIALS AND METHODS

In this section, we will introduce the key steps involved in addressing the data quality problem through a generative approach. We will discuss the dataset preparation process, including the construction of source and target sets based on membrane quality and cell types, as well as the integration of manual annotation for improved categorization. Furthermore, we will detail the training of CycleGAN models with generators and discriminators to learn the mapping between the source and target sets, ensuring the generation of realistic and enhanced membranes. Based on our goals, we design three different experimental settings to build three generation models, where two of the generators aim to separately impute tumor cell membranes and immune cell membranes given only nuclei signals and the third model will generate the whole tumor membranes for cells with weak and partial membranes. Finally, we will explain how these trained models can be applied during testing to generate complete membranes for immune cells or tumor cells with weak or no membrane staining, ultimately improving the accuracy and reliability of cell segmentation.

### 2.1 Data acquisition and image processing

In our study, we utilize the publicly available CosMx dataset [10] as our primary dataset. This dataset encompasses a comprehensive collection of multiplexed immunofluorescence images and it is chosen due to its wide variety of cell types and staining conditions, allowing us to evaluate the performance of our generative approach across diverse scenarios.

There are totally eight formalin-fixed, paraffin-embedded (FFPE) non-small cell lung cancer (NSCLC) tissue samples, 120 images or fields of view (FOV) consisting of around 390,000 cells, which are obtained using the spatial molecular imaging technique from CosMx platform [10]. Four samples “Lung5_Rep1”, “Lung5_Rep2”, “Lung5_Rep3”, and “Lung6” are utilized for training and evaluating our methods. These samples were specifically selected for their suitability in containing a substantial number of cells with relatively well-integrated membrane signals. This selection ensures that the membranes generated by our approach meet a high standard of quality. These samples are obtained from the same patient, while “Lung6” is obtained from a separate patient. Specifically, “Lung5_Rep1”, “Lung5_Rep2”, “Lung5_Rep3”, and “Lung6”, contain 32, 30, 32, and 30 FOVs, each with 98,002, 105,800, 97,809, and 89,975 cells, respectively. Each sample comprises 18 distinct cell types, including one identified as cancerous. The sample images are stained with both nuclear and membrane markers (DAPI, CD298, PanCK and CD3).

Different membrane channels are first combined and then normalized to the range of the nuclear channel. Next, the nuclear channel is subtracted from the combined membrane channel to enhance the signal contrast. These processed channels are then used to create images, which can be further used to segment nuclear and membrane and ensure an appropriate cell boundary definition by combining both. These images are high-resolution, and each image (FOV) with a size of 5,472 pixels × 3,648 pixels, 0.18 *µm* per pixel contains around 3,000 densely gathered cells, making it challenging to train generation models to focus on specific cell types and membrane quality. Therefore, each image were firstly cropped into a set of patches with size 456 × 456, and the background patches were then manually filtered to ensure each patch contains 10 ∼ 50 cells.

To achieve our objective, we construct specific source-domain patch sets and target-domain patch sets for training each model and enabling the transfer from the source style to the target style, as illustrated in Fig. 3. The source-domain patches consist of two distinct sets: immune/tumor cells without membranes and tumor cells with weak membrane staining. On the other hand, the target-domain patches comprise sets of immune cells with well-defined membranes and tumor cells with well-defined membranes. These different source and target sets form various pairs of training sets tailored for each of the three models in our framework. This approach facilitates the training process and enables the models to learn the transformation from the source-style, with low-quality or absent membranes, to the target-style, featuring high-quality membranes, separately for immune and tumor cells.

**Figure 3:**
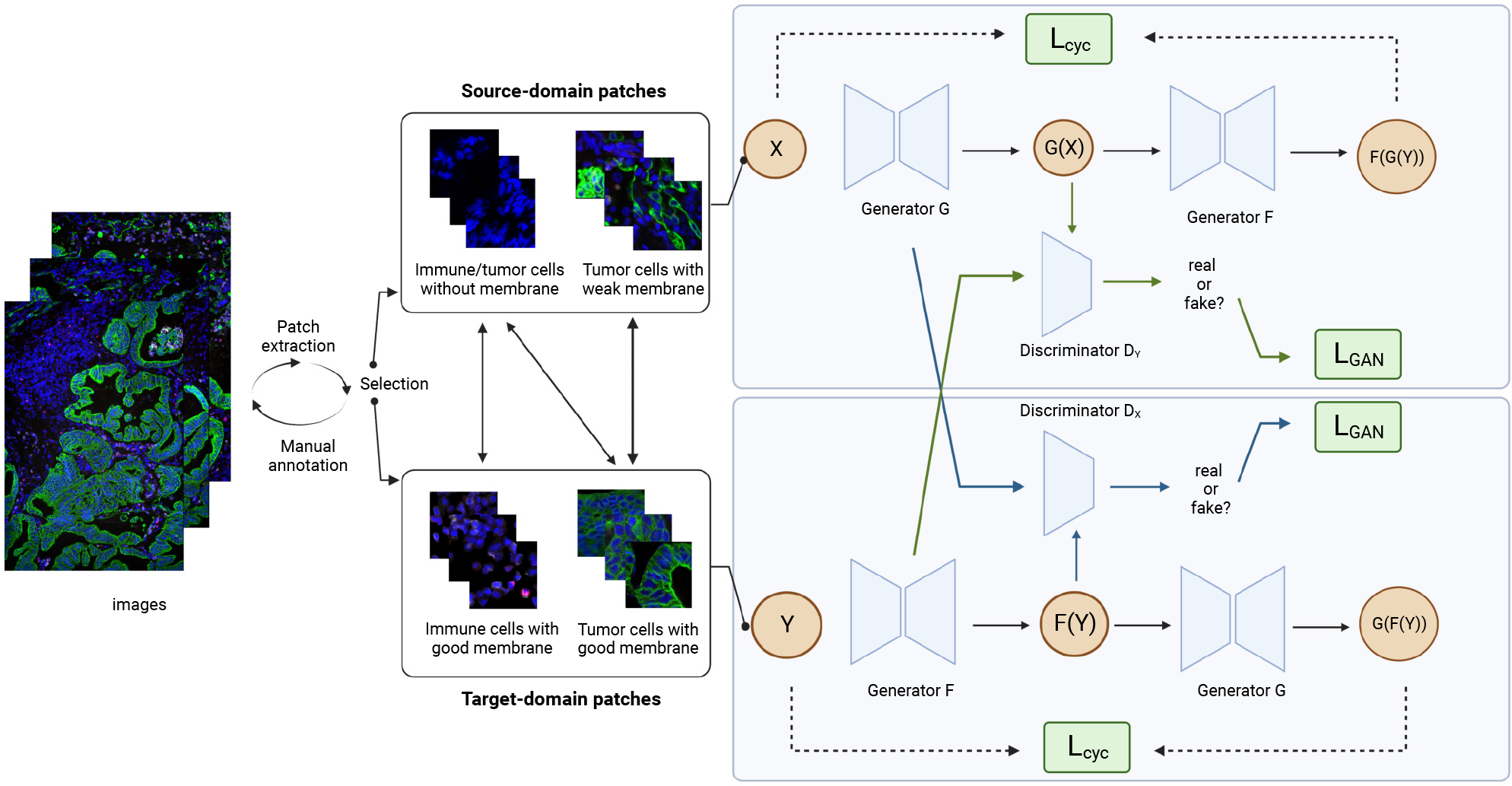
The schematic overview of the MemGAN framework. Multiple source-domain and target-domain sets are manually constructed, forming the basis for generation models in three different combinations: nuclei to immune cells, nuclei to tumor cells, and weak membrane of tumor cells to good membrane. All three models adhere to the same training pattern, employing Cyclegan as illustrated in the right part of the diagram.

Since the cells are randomly distributed, each patch may contain both tumor and immune cells. Therefore, all the patches were first manually divided into two sets: one with the most tumor cells in each patch and the other with the most immune cells. To ensure that the model can synthesize the target membranes without noisy information, five well-trained annotators manually mask the tumor cells and immune cells with both nuclei and membrane areas separately based on the fluorescent staining so that the two sets separately contain only tumor cells and only immune cells. To generate high-quality membranes, cells with incomplete or noisy staining were also filtered out from the two sets. These two filtered sets, denoted as 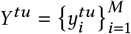 and 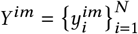, can be utilized as the target datasets for membrane generation of tumor cells and immune cells, where they follow the data distribution *y*^tu^ ∼ *p*_data_ (*y*^tu^) and *y*^im^ ∼ *p*_data_ (*y*^im^). As the aim of the first two models is to impute membranes for cells with only nuclei signals, we built the corresponding source datasets, 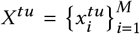 and 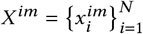, using the nuclei channels of *Y*^tu^ and *Y*^im^, where *x*^tu^ ∼ *p*_data_ (*x*^tu^) and *x*^im^ ∼ *p*_data_ (*x*^im^). For the third model, the source dataset 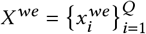 was constructed from a manual selection of patches with partial and weak membranes, so this model could be trained to impute the missing membrane signals for the incomplete cells in *X* ^*we*^.

### 2.2 Membrane generation with Mem-GAN

Generative Adversarial Network (GAN) is a widely used deep neural network for generation tasks, such as image synthesis [2, 25, 35, 37], text-to-speech [14, 20, 28, 38], and video prediction [8, 11, 19]. The training process is a “min-max” game between a generator and a discriminator, where the generator try to produce realistic samples and the discriminator aims to distinguish between the generated samples and the real training data. Based on GAN, there are advances on image-to-image translation tasks, which transfers the style of one image domain to another domain. Unlike Pix2pix [12] which is limited to paired training dataset, CycleGAN [39] has been a powerful tool to map images from a source domain and a target domain with unpaired data. Since the task of cell membrane generation is much more challenging, where the source data *X* and target *Y* are mostly unpaired, we adopted the CycleGAN as the architecture of each generator to map the image distribution between the source *X* and target *Y*.

As shown in the right part of Figure. 3, the whole process includes two mapping functions *G* : *X* → *Y* and *F* : *Y* → *X*. For the first translation model of generating membranes from nuclei for tumor cells, the two generators *G* : *X*^*tu*^ → *Y* ^*tu*^ and *F* : *Y* ^*tu*^ → *X*^*tu*^ are used for translating from nuclei to membrane and from membrane to nuclei, respectively. At the same time, we have two discriminators 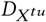 and 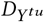, which are trained to distinguish between cells {*G* (*x*^*tu*^) }, { *F* (*y*^*tu*^)} generated by *G, F* and the real cells { *y*^*tu*^ }, {*x*^*tu*^ *}* from {*X*^*tu*^ } and {*Y*^*tu*^}. These process can be optimized via the following adversarial loss formulation:

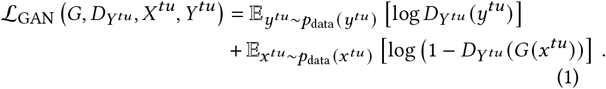

The loss function 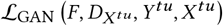 for the mapping *F* is similar.

To map each individual cell to its corresponding membrane rather than a random mapping to the target domain *Y* ^*tu*^, just to fit the distribution *p*_data_ (*y*^*tu*^), a cycle consistency loss is introduced to reconstruct *G* (*x*^*tu*^) back to *x*^*tu*^ using mean squared error (MSE), and bring *F* (*y*^*tu*^) back to *y*^*tu*^ as well:

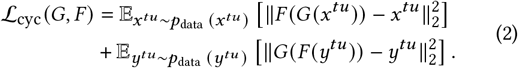

The overall loss function is defined as:

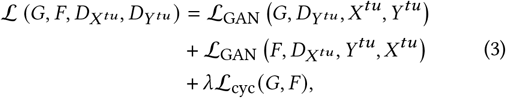

where *Ji* is set as 10 to weight the losses. Therefore, the ultimate goal of the overall training procedure is the following optimization:

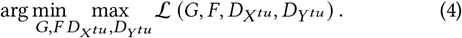

This translation process is particularly trained for tumor cells, which enhances the similarity between the distribution of tumor cells with synthetic membranes and real cells.

Similarly, the second model includes mappings *G* : *X*^*im*^ → *Y* ^*im*^ and *F* : *Y* ^*im*^ → *X*^*im*^, and the third model tries to map *G* : *X* ^*we*^ → *Y* ^*tu*^ and *F* : *Y* ^*tu*^ → *X* ^*we*^, with their corresponding discriminators. They have the similar optimization process.

Once the generative models have been trained, they can individually produce missing membranes by taking nuclei channels as the input or images with nuclei and weak membranes. In the post-processing step, we employ a reconstruction process to merge the original nuclei channels with the newly generated membrane channels, thereby preserving the genuine nuclei signals without any loss.

### 2.3 Cell segmentation from generated cells

Given the generated fluorescent microscopy, segmentation can be performed to identify and isolate individual cells or other structures within the image. In our segmentation process, we adopt Cellpose [33, 34] as a strong segmentation tool to conduct cell segmentation. This U-Net-based model has been trained on large manually annotated images and can perform the segmentation of cells with variable shapes, sizes, and multiple channels in fluorescence microscopy images. Therefore, with our enhanced membranes, Cellpose is expected to generate more accurate boundaries for further single-cell analysis. Moreover, since the accuracy on single-cell segmentation is largely based on the integrity and quality of the membrane signals, cell segmentation can also serve as an effective way to evaluate the quality of the generated cells.

## 3 RESULTS

In this section, we assess the performance of our three generation models using the CosMx dataset [10]. We visualize the generated cells and indirectly evaluate the quality of the generated membranes by comparing the segmentation results both qualitatively and quantitatively. Given the lack of “ground truth” with respect to cells with weak membranes, we incorporate human evaluation to appraise the generation performance and highlight the improvements in cell segmentation.

### 3.1 Experimental settings

All the experiments were implemented in Pytorch and trained on Nvidia RTX A6000 GPU. The Adam optimizer was employed with an initial learning rate of 0.0002 for the initial 100 epochs. Additionally, a batchsize was set as 1 during training and testing phase. To ensure gradual convergence, we linearly decayed the learning rate to zero every 50 epochs over the remaining 100 epochs of training. We randomly split the cropped patches into training and testing sets by a ratio of 4:1. The whole model was trained in an end-to-end manner.

### 3.2 Evaluation of generated tumor cell membrane

To assess the effectiveness of the proposed membrane generation model on tumor cells based solely on nuclei, we perform a comparative analysis between the original images containing both nuclei and membrane signals and the post-processed images generated by our models.

The visual comparison of the two cases is depicted in Figure. 4. The first column of each case represents the original image, displaying clearly visible and integrated tumor cell membranes. Mem-GAN, with only the nuclei channels as input, generates similar membrane signals compared to the ground-truth images while maintaining the original integrated nuclei. Notably, the generated tumor cells exhibit distinct membrane boundaries that are welldefined.

**Figure 4:**
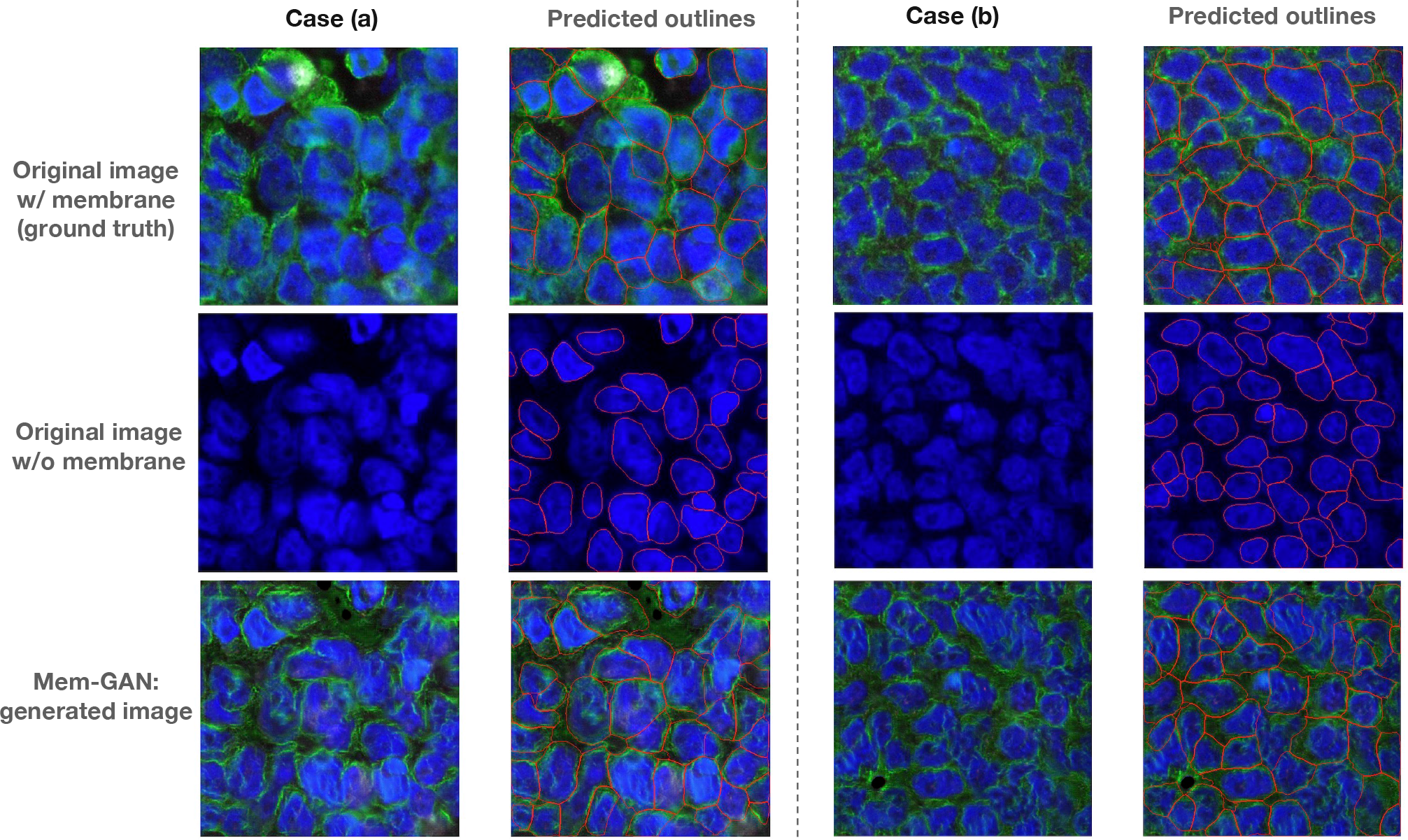
Visual comparison of original tumor cells and tumor cells generated by our method. The first and second columns of Case (a) and Case (b) display the images and the corresponding cell boundaries predicted by Cellpose [34], respectively.

The ultimate objective of Mem-GAN is to produce complete and integrated membrane markers for single cells, which can result in more precise and accurate identification and segmentation of individual cells in fluorescent microscopy images. In Figure. 4, the second column of each case illustrates the segmentation results of Cellpose based on original images generated images from Mem-GAN. When given only the nuclei channels, Cellpose recognizes only the boundaries of the nuclei, and some cells are missed. By employing our generated membranes, Cellpose can detect most of the missing cells, and the “fake” membranes can be effectively utilized to determine the approximately real boundaries. Table 1 presents the quantitative evaluation of cell segmentation by computing the intersection-over-union (IoU) score between the best-matched masks and the corresponding ground truth masks. These masks are generated from continuous pixel values, where each pixel is assigned to either a specific cell or the background. The mask IoU results are shown for patches from three different samples, with the segmentation boundaries of the original image with membranes used as the ground truth. The results indicate that Mem-GAN significantly improves the IoU score by up to 0.068 for sample “Lung6”, with an average improvement of 10% compared to the original images without membranes. Therefore, with the improved capturing of single-cell structures, it can further enhance downstream analysis and enables the investigation of cell heterogeneity, the identification of rare cell types, and the study of dynamic cellular processes.

**Table 1:** Quantitative results of cell segmentation using Mem-GAN for membrane generation.

|  | Original image w/o membrane | MEM-GAN |
| --- | --- | --- |
| Lung5_Rep1 | 0.2855 | 0.2943 |
| Lung5_Rep2 | 0.2956 | 0.3133 |
| Lung5_Rep3 | 0.2501 | 0.2743 |
| Lung6 | 0.2839 | 0.3516 |
| Average | 0.2788 | 0.3084 |

Hence, in situations where the membrane markers are missed, our Mem-GAN can effectively generate genuine membranes for each cell. The authenticity of our generated membranes is not only verified through the visual comparison with the original ground truth images, but also by the downstream single-cell segmentation based on the nuclei and membrane signals.

### 3.3 Evaluation of generated immune cell membrane

Similar to evaluating the tumor membrane generation, we assess the second generation model of generating membranes for immune cells. Compared with tumor cells which exhibit heterogeneous expression of markers, immune cells generally have a round shape with a large nucleus and a more uniform appearance. In our images, immune cells exhibit weaker membrane signals due to the potential interference from tumor cell markers during the image acquisition process. This may lead to signal quenching, causing the immune cells to have faint or incomplete membrane signals. These weak signals can be considered as inevitable noise in the models. Despite this, as demonstrated in Figure. 5, the first column of the two cases shows that our model can still generate marker signals when provided only with nuclei channels as input. Interestingly, when compared to the original images, the model even aids in amplifying the signals of certain cells.

**Figure 5:**
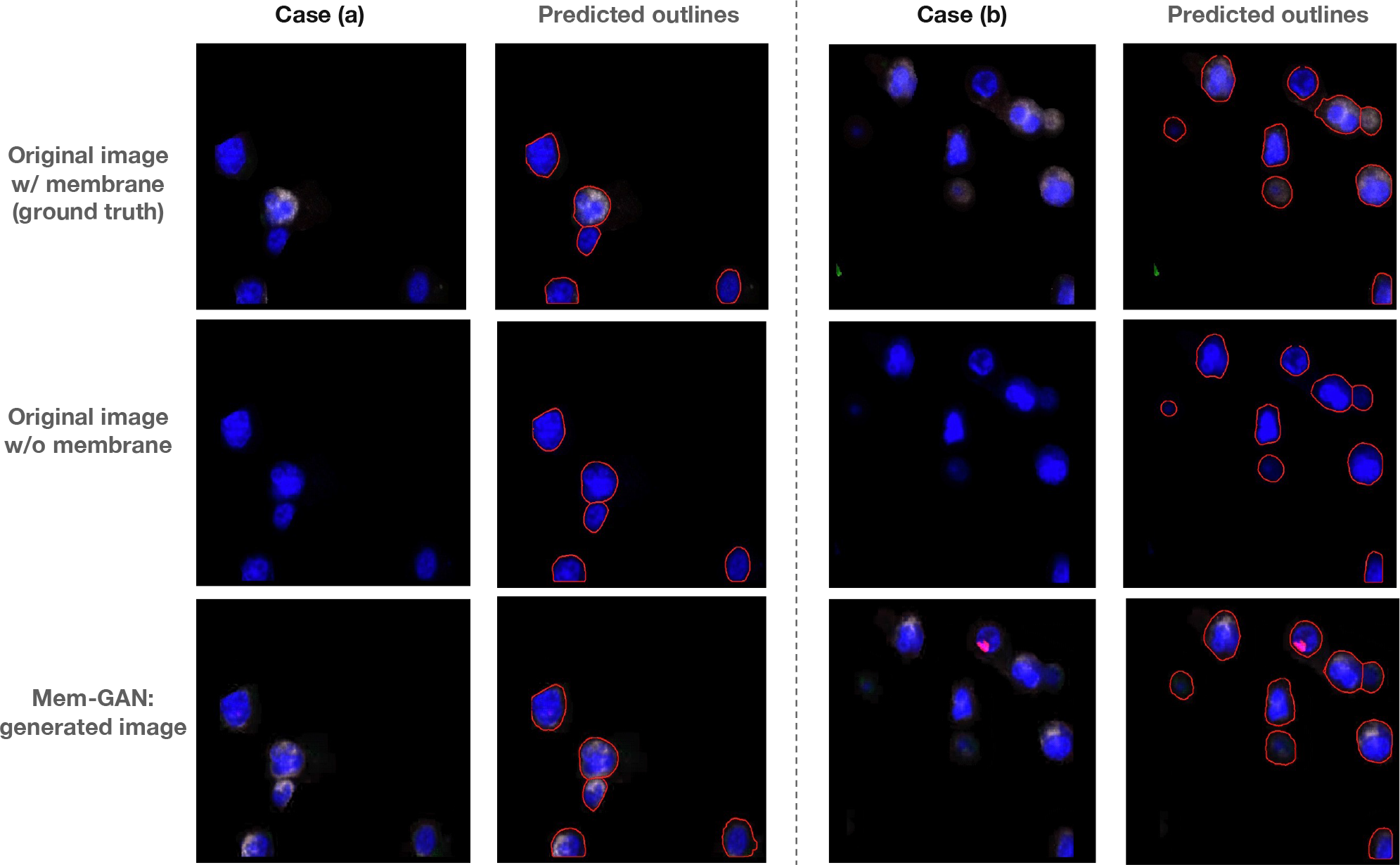
Visual comparison of original immune cells and immune cells generated by our method. The first and second columns of Case (a) and Case (b) display the images and the corresponding cell boundaries predicted by Cellpose [34], respectively.

Meanwhile, immune cells may be more sparsely distributed, especially when infiltrating the tumor tissue, making them easier to segment compared with tumor cells. As shown in the second column, the pseudo membranes in our generated images can be advantageous for segmentation, enabling more accurate identification of individual cells.

### 3.4 Performance on weak membrane generation

The first two models primarily serve to confirm the accuracy of the generated results when utilizing solely nuclei channels, given that we have access to the original images as a point of reference. Based on high similarities between the generated and actual membranes, our third model addresses a more complicated, yet commonplace, situation where the membranes are partially absent or weak.

Figure. 6 depicts four cases that demonstrate the significant enhancement of tumor cell membrane achieved by Mem-GAN. Additionally, it displays the corresponding segmentation boundaries generated by Cellpose and the spatial gradient map derived from the simulated diffusion process within Cellpose. These four cases demonstrate several distinct scenarios involving weak membrane signals. For instance, Case (a) depicts numerous cells with partial or indistinct membrane signals, which are effectively repaired by Mem-GAN based on the existing signals. Case (b) represents a more challenging scenario than Case (a), as it involves cells without any detectable membrane signals. Despite this difficulty, Mem-GAN can generate relatively authentic membranes for these cells while retaining the essential initial signals. In these two cases, Cellpose struggles to identify the cells with incomplete or missing membranes. However, the predicted outlines suggest that the generated membranes significantly aid the segmentation model in capturing the cell boundaries. Case (c) and Case (d) illustrate the scenario where some existing membrane markers are unreliable and inaccurate, such as membranes without nuclei or membranes that are free-floating. These challenges significantly increase the difficulty of the membrane generation task. It is worth noting that Mem-GAN may be misled by these “fake” membranes, as the model aims to generate more signals based on the available ones. Despite this limitation, it is evident in the segmentation results that our model is still able to generate membranes that are beneficial for the segmentation process, as depicted in the second row of Figure. 6.

**Figure 6:**
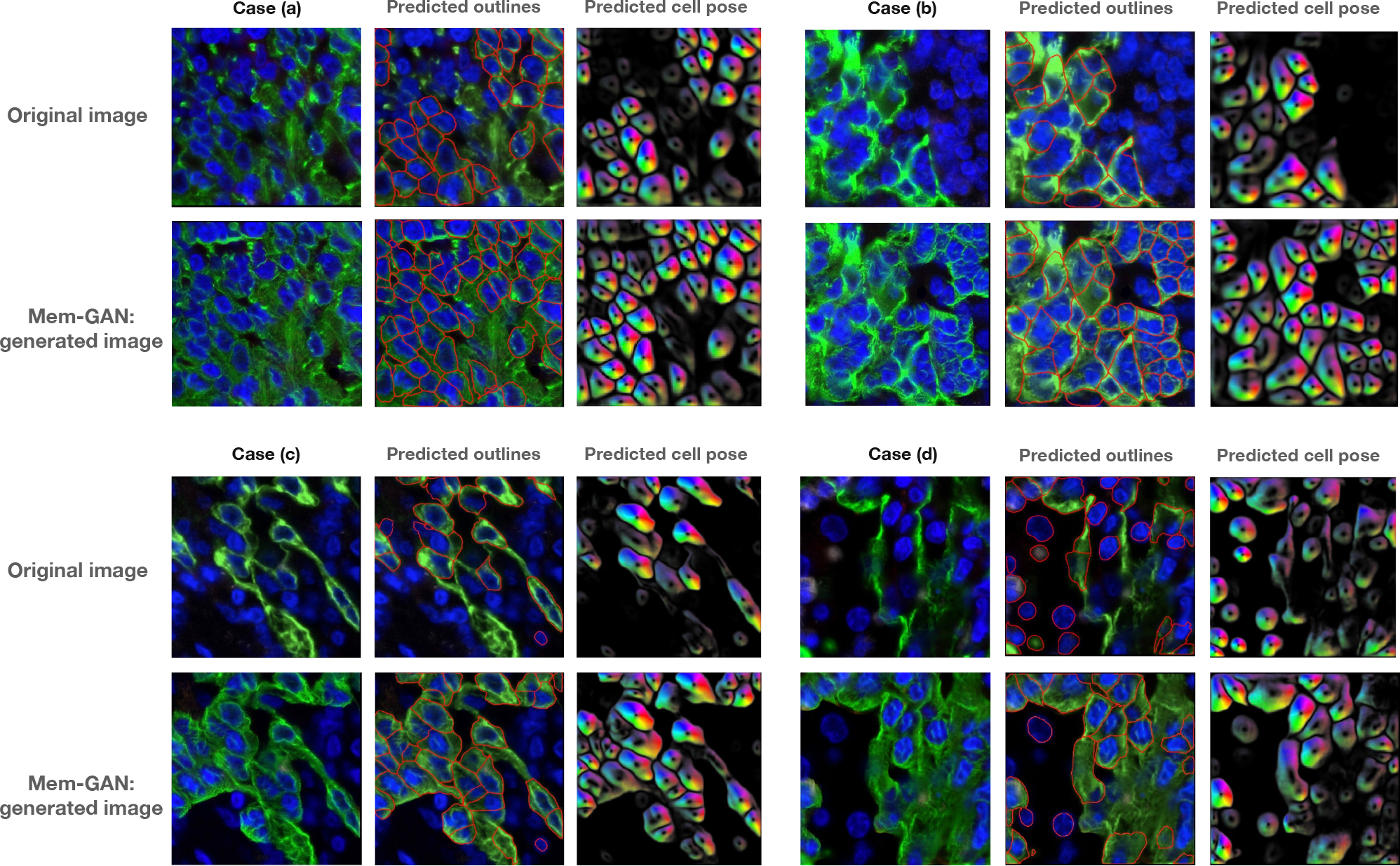
Qualitative results for weak membrane generation. The original images in the dataset exhibit low staining quality, specifically weak membrane signals. Our generated images enhance and complement the lost signals, resulting in improved cell segmentation, as demonstrated in the second and third columns for each case.

### 3.5 Human evaluation on weak membrane generation

Due to the absence of ground truth for cells with originally weak membranes, objectively assessing the authenticity of the generated membranes becomes challenging. Therefore, we adopt a manual evaluation process to assess the quality of the generated membranes and their impact on cell segmentation. This evaluation involves five experienced PhD students with domain expertise who manually inspect and judge the generated membranes and the resulting segmentation outcomes. Notably, these evaluators are unaware of whether the images they evaluate are original or generated by Mem-GAN, ensuring an unbiased assessment process. While subjective, this evaluation provides valuable insights into the quality and effectiveness of the generated membranes in improving cell segmentation. Each student is tasked with evaluating a total of 40 image pairs, consisting of 20 pairs from “Lung5_Rep1” and 20 pairs from “Lung5_Rep2”. In each image pair, one image undergoes cell segmentation using the original membrane staining, while the other image undergoes cell segmentation using our generated membrane staining through Mem-GAN. The evaluation metrics employed in our study include “Membrane Completeness”, “Membrane Quality”, and “Cell Segmentation”. Each metric is categorized into five levels, with corresponding scores ranging from 1 to 5. The scoring scale is defined as follows: Score 1 represents “Very poor”, Score 2 represents “Poor”, Score 3 represents “Acceptable”, Score 4 represents “Good”, and Score 5 represents “Very Good”. For the assessment of Membrane Completeness, the students are instructed to assign scores based on the percentage of cells exhibiting membrane staining. A score of 5 is assigned if over 90% of cells show membrane staining. Scores of 4, 3, 2, and 1 are assigned for percentages of 80%-

90%, 60%-80%, 40%-60%, and less than 40%, respectively. Similarly, for the evaluation of Membrane Quality, the students consider the ratio of cells with well-defined, separable, and clear membranes. The corresponding score is assigned based on the defined ranges above. In the case of Cell Segmentation, the students assess the ratio of cells correctly identified and segmented. The score for this metric is determined based on the observed ratio of cells to the specified ranges above.

As observed in Table 2, it is evident that both in “Lung5_Rep1” and “Lung5_Rep2”, the original images exhibit incomplete membrane staining with relatively low membrane quality, scoring around 2. Additionally, the cell segmentation score of 2 indicates that only approximately half of the cells are successfully identified. However, by employing Mem-GAN, the generated pseudo membrane significantly aids in membrane generation for cells lacking proper staining. As a result, there is a notable improvement across all three metrics. Moreover, our algorithm’s pseudo membrane staining greatly enhances the performance of cell segmentation.

**Table 2:** Human evaluation in terms of Membrane Completeness, Membrane Quality and Cell Segmentation. Each evaluation metric will be categorized into five levels with corresponding scores ranging from 1 to 5. The higher values indicate better performance. Each score rating for each metric is accompanied by its specific definition, which can be found in the detailed explanation provided in the text.

|  | Image | Membrane Completeness | Membrane Quality | Cell Seg |
| --- | --- | --- | --- | --- |
| Lung5_Rep1 | Original | 2.25 | 2.35 | 2.4 |
|  | MEM-GAN | 3.4 | 3.15 | 3.4 |
| Lung5_Rep2 | Original | 1.7 | 2.2 | 2.3 |
|  | MEM-GAN | 3.45 | 3.1 | 3.35 |
| Average | Original | 1.975 | 2.275 | 2.35 |
|  | MEM-GAN | 3.425 | 3.125 | 3.375 |

## 4 DISCUSSION

The reliability of membrane markers for cell segmentation can be compromised by several data quality factors, such as staining artifacts, imaging noise, and limited microscope resolution. These factors result in missing or unclear membrane staining, leading to uneven and low-quality staining. Traditional and advanced deep learning-based cell segmentation methods rely significantly on the staining quality of input images, and as a result, they may encounter limitations when dealing with complex data featuring diverse cell shapes, high cellular density, and unclear cell boundaries.

Instead of improving the segmentation algorithms, in this work, we propose a new perspective that addresses the inadequate staining problem at its essence. Our approach involves generating missing or unclear membrane markers using generative models, which aims to improve the quality of the stained images and enable more precise identification and segmentation of individual cells. This method provides an effective solution to the challenge of inadequate staining and it has the potential to significantly assist single cell analysis in various downstream applications.

Despite its effectiveness in generating authentic pseudo membranes for tumor and immune cells, our current model encounters limitations when confronted with out-of-distribution tissues and cells displaying complex morphologies. In such cases, additional training data from diverse datasets may be required to adapt to new membrane staining patterns. Nevertheless, it’s important to note that Mem-GAN serves as a promising proof-of-concept for improving staining quality through the application of generative models and deep neural networks.

A prospective avenue for future research involves the development of a single cell-based image classifier capable of distinguishing different cell types, such as immune cells or tumor cells, prior to applying Mem-GAN. By doing so, the corresponding models can be employed to generate appropriate pseudo membranes tailored to each cell type. However, even in scenarios where cell type differentiation proves challenging, there is still potential value in utilizing the generated tumor cell membranes for immune cells to aid in cell boundary identification. In the context of segmentation, the ultimate goal remains the delineation of complete cell boundaries. Therefore, generating pseudo membranes for immune cells using the tumor cell generator can still contribute to improved segmentation results. These generated membranes offer valuable additional information that assists Cellpose or any other segmentation tools in recognizing and segmenting cells, ultimately enhancing segmentation outcomes. As part of future work, the exploration of more advanced generative models, such as diffusion models, holds promise in further enhancing the capabilities of Mem-GAN.

